# Emergence of Human Amygdala Functional Networks: 3 Months to 5 Years of Age

**DOI:** 10.1101/261347

**Authors:** L.J. Gabard-Durnam, J. O’Muircheartaigh, H. Dirks, D.C. Dean, N. Tottenham, S. Deoni

## Abstract

Although the amygdala’s role in shaping social behavior is especially important during early post-natal development, very little is known of human amygdala functional development before childhood. To address this important gap, this study used resting-state fMRI to examine early functional network development of the amygdala and its subregions in 80 participants from 3-months to 5-years of age. Whole brain functional connectivity with the whole amygdala and its laterobasal and superficial nuclear groups were largely similar to those seen in older children and adults, and functional distinctions between subregion networks exist already. These patterns suggest many amygdala functional circuits are intact from infancy, especially those that are part of larger motor, visual, auditory and subcortical (basal ganglia especially) networks. Notably, these observed robust amygdala functional networks in infancy precede reports to date of elicited amygdala reactivity in development. Developmental changes in connectivity were observed between the laterobasal nucleus and bilateral ventral temporal and motor cortex as well as between the superficial nuclei and medial thalamus, occipital cortex and a different region of motor cortex. These results show amygdala-subcortical and sensory-cortex connectivity begins refinement prior to childhood, though connectivity changes with associative and frontal cortical areas, seen after early childhood, were not evident in this age range. These findings represent early steps in understanding amygdala network dynamics across infancy through early childhood, an important period of emotional and cognitive development.

## Introduction

The amygdala plays a central role in the processing and production of emotional and social behavior [Adolphs and Spezio, 2006; Ochsner et al., 2012; Phillips et al., 2003], both through its own activity as well as its dense anatomical and functional connectivity to the rest of the brain [Kim et al., 2011; Phelps and LeDoux, 2005; Young et al., 1994]. Rich animal and human literatures examining the effect of both lesions and stress on the amygdala across the lifespan suggest that this region’s role in shaping emotion and social behavior is especially important during early post-natal development, with consequences for affective behavior lasting throughout life [Adolphs et al., 1994; Adolphs et al., 2002; Bauman et al., 2004; Bliss-Moreau et al., 2011; Bliss-Moreau et al., 2013; Graham et al., 2015a; Kazama et al., 2012; Prather et al., 2001; Raper et al., 2013; Shaw et al., 2004]. Due to these associations, clarifying the role of the amygdala and its networks during early neurodevelopment is imperative to understanding the ontogeny of affective phenotypes over typical and atypical development [Graham et al., 2015a; Roy et al., 2013; Shen et al., 2016; Tottenham and Sheridan, 2009].

Therefore, work in humans has focused on examining the developing amygdala. These studies have revealed a robust functional responsiveness to emotional stimuli by childhood (4-10 years) [Dehaene-Lambertz et al., 2010; Gee et al., 2013; Hare et al., 2008; Perlman and Pelphrey, 2011; Swartz et al., 2014; Thomas et al., 2001]. In the absence of stimuli, variations in the amygdala’s ongoing functional connectivity at just 1 month of age have been associated with affective and cognitive profiles 5 months later [Graham et al., 2015a]. By childhood, the amygdala exhibits functional connectivity with diffuse brain regions (in particular subcortical and temporal cortical regions, Shen et al., 2016). Functional connections with other cortical regions, especially the prefrontal cortex, continue to develop from childhood into adulthood [Alarcón et al., 2015; Gabard-Durnam et al., 2014; Qin et al., 2012].

An emerging focus in the study of the developing human amygdala is on understanding the early functional organization of the amygdala’s anatomical subregions. An expansive animal literature has established how these subregions serve different functional roles in maturity. Specifically, the laterobasal nuclei are central to aversive and appetitive valuation through cortical and subcortical connections [Ghashghaei and Barbas, 2002; Killcross et al., 1997; Pessoa and Adolphs, 2010]. In contrast, the superficial nucleus processes olfactory and social information through connectivity with the insula and piriform cortex [Bzdok et al., 2013; Heimer and Van Hoesen, 2006; McDonald, 1998]. Similarly, each of these subregions in humans has been shown to serve different affective functions through distinct activity and network connectivity [Ball et al., 2007; Davis et al., 2010; Etkin et al., 2009; Frühholz and Grandjean, 2013; Price, 2003; Roy et al., 2009, Kerestes et al., 2017]. Preliminary developmental studies in older children (4-10 years) suggest that progressive functional segregation between these subregions’ cortical networks occurs throughout childhood and adolescence [Gabard-Durnam et al., 2014; Qin et al., 2012; Roy et al., 2013]. However, very little is known of the functional development of the human amygdala or its constituent subregions across infancy and early childhood. This represents a significant gap in our understanding of this early-developing structure.

Therefore, the current study assessed amygdala networks from infancy through early childhood using resting-state functional magnetic resonance imaging (RS-fMRI). Specifically we investigated early network development of the whole amygdala as well as two of the amygdala’s major subdivisions, the laterobasal and superficial subregions, across infancy and early childhood. RS-fMRI provides a robust measure of functional network composition indexing the maintenance and stability of functional connections [Larson-Prior et al., 2009; Pizoli et al., 2011; Raichle, 2010]. Additionally, RS-fMRI can facilitate the identification of developmental changes in functional networks as they emerge without task design confounds or confounds due to differences in cognitive functioning during the period of interest [Graham et al., 2015b; Uddin et al., 2010]. Thus, RS-fMRI approaches enabled this study to begin addressing the paucity of understanding about human amygdala networks during early development.

## Methods

### Participants

#### Paediatric Data

Informed consent was obtained from each child’s parent or guardian in accordance to ethics approval from the Brown University Institutional Review Board. The participants from this study were enrolled as part of a larger longitudinal study of typical brain development [Dean et al., 2014; Deoni et al., 2012] and were screened at enrollment to exclude major risk factors for developmental delay or psychopathology. Inclusion criteria included: uncomplicated single birth between 37 and 42 weeks gestation; no reported in utero exposure to alcohol or illicit drugs; and no first degree familial or personal history of major psychiatric or neurological illness as reported by caregivers. All data used in this analysis were acquired while the infant or child was sleeping naturally using previously reported imaging protocols and techniques [Dean et al., 2014]. All infants and children were asleep for 15-20 minutes before entering the MRI machine.

A total of 159 datasets were initially collected from 133 individual children between 3 and 54 months of age. Of the initial 159 datasets, 50 datasets were excluded based on an initial screening of motion characteristics of the functional data, (see Functional MRI section below). A further 8 were excluded due to low quality of the structural scan. Finally, for children who were scanned on more than one occasion, only their later dataset was included, leading to the exclusion of a further 21 datasets. The remaining 80 participants, between 84 and 1682 days of age corrected to a 40 week gestation, contributed usable data for these analyses (see Table 1 for sample characteristics, stratified by age group for visualization only, these categorical groups were not used in analysis). Of this sample of 80, self-reported ethnicity was collected for 70 subjects. Of these, parents or guardians reported the ethnicity of their children as European American (40/70, 57.1%), African American (8/70, 11.4%), Asian American (1/70, 1.4%), mixed background (17/70, 24.3%) or other (4/70, 5.8%). Of these, 15/70 (21.4%) identified as Hispanic.

**Table 1:** Sample characteristics for participants contributing usable data.

| Age Group | N | Sex Ratio (M:F) | Average Age (days) |
| --- | --- | --- | --- |
| 0-11.9 months Old | 25 | 13:12 | 160 |
| 1-1.9 Years Old | 21 | 12:9 | 565 |
| 2-2.9 Years Old | 13 | 7:6 | 939 |
| 3-5 Years Old | 21 | 12:9 | 1374 |

#### Adult data

We also investigated an adult dataset for comparison. Data were accessed from the International Neuroimaging Data-sharing Initiative ( http://fcon_1000.projects.nitrc.org/) and we used the NKI-Rockland dataset (Nooner et al, 2012), a community based sample of children and adults aged 6-85 years. The initial sample size available was 209, but as in our previous work (O’Muircheartaigh et al, 2015), this was reduced to 102 after restricting the sample to adults aged 18-65 years (excluding 71) and only including datasets that were both complete and passed strict quality control criteria (excluding 36, see functional MRI section below). The remaining 102 participants (40 female) had a mean age of 33.9 years (standard deviation 12.9).

### Structural MRI

#### Paediatric Data

MRI data were acquired on a single Siemens _3_T Tim Trio equipped with a 12-channel radio-frequency head array. The T_1_ weighted SPoiled GRadient echo (SPGR) structural image used here was acquired as part of the mcDESPOT sequence (multi-component driven equilibrium single pulse observation of T1 and T2, Deoni et al., 2008, 2012). This sequence is designed to enable calculation of the signal in water attributable to different tissue populations, but here we used only the high flip-angle T_1_ weighted image for registration and morphometry. The acquisition parameters for this sequence were optimized according to age group (Deoni et al., 2012) and expected head size and images were acquired sagittally (www.cdc.gov/growthcharts/clinical_charts.htm). This also means that the acquisition parameters (for only this structural scan) varied with age (see **Table 2**).

**Table 2:**
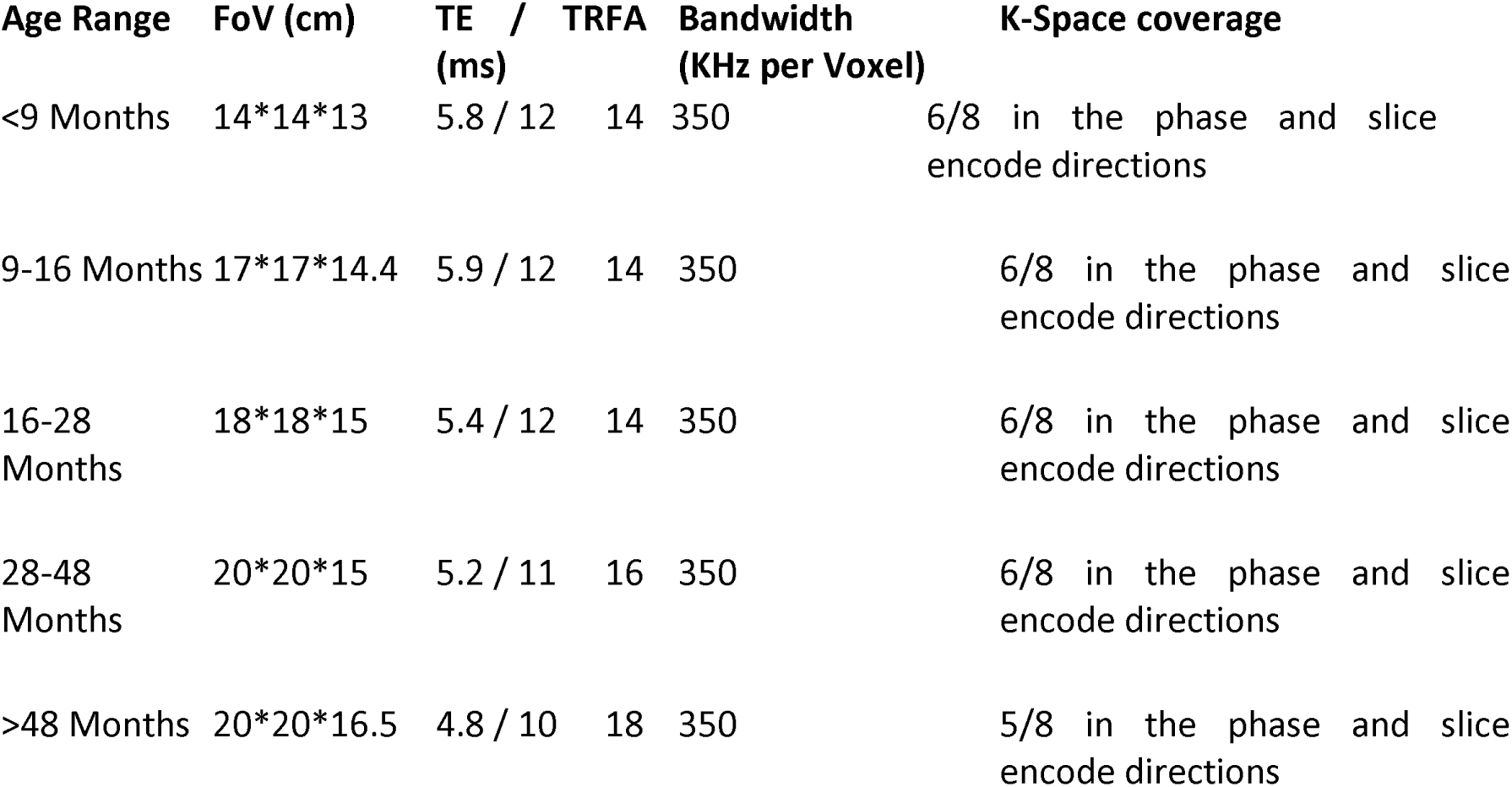
Acquisition parameters for the T1 weighted SPGR data for each age group.

Each T_1_ image was first corrected for intensity non-uniformities using the N_4_BiasFieldCorrection tool [Tustison et al., 2010] and then stripped of non-brain tissue using the Brain Extraction Tool [Smith, 2002]. The resulting image was non-linearly registered to a series of age-specific T_1_ weighted template images derived from an independent population of infants and toddlers who were scanned with the same sequence [Deoni et al., 2012], and then registered to an overall average paediatric brain template using the Advanced Normalization Tools [Avants et al., 2008]. In this way, all images were normalized to the same space.

#### Adult data

All adult data were acquired on a different Siemens 3T Siemens Tim Trio. T_1_-weighted images were collected using a magnetization prepared rapid gradient echo (MPRAGE) sequence with a TR/TE of 2500 ms/3.5 ms, inversion time of 1200 ms, flip angle of 8°, and a field of view of 256 mm × 256 mm over 192 slices resulting in an isotropic resolution of 1 mm. For every subject, the resulting image was non-linearly aligned to the MNI template (adult) template using fnirt (part of the FSL package; Andersson et al, 2010).

### Functional MRI

#### Paediatric Data

A gradient echo-planar imaging sequence was used for functional MRI data acquisition with the following parameters for all participants in the study: repetition time=2.4s, echo time=34ms, flip angle=80°, 32 interleaved 3.6mm slices acquired at an orientation parallel to the anterior-posterior commissure line, in-place resolution of 2.97mm, and a GRAPPA acceleration of 2. A total of 132 volumes were initially collected but the first 4 were discarded to allow magnetization to reach equilibrium leading to just over 5 minutes of data (307.2 seconds) per subject. Each resulting time-series was corrected for inter-scan motion using MCFLIRT (FMRIB Software Library [FSL], Oxford, United Kingdom [Smith et al., 2004] and had the middle functional scan rigidly registered to its T_1_ weighted image.

Following this preprocessing, basic motion characteristics were examined. As this analysis focuses on an area near the edge of the brain and surrounded by vasculature and at risk of susceptibility signal dropout, the criteria used to exclude / include datasets were strict. For each time-series, the number of volumes with >0.2mm scan-to-scan motion was recorded, consistent with prior infant studies of resting-state connectivity [e.g. Gao et al., 2014; Graham et al., 2015b]. Any time-series with >12 (i.e. 10%) of scans with these motion spikes (42 datasets) or with absolute or relative motion over the course of the time-series of >2mm (8 datasets) were discarded from further analysis. The remaining datasets were corrected for structured noise associated with motion artifact using the AROMA package [Pruim et al., 2015] (v0.3b available at https://github.com/rhr-pruim/ICA-AROMA). AROMA is an independent component analysis (ICA) based method that automatically classifies and removes components identified as noise according to a combination the correlation of each components timeseries with motion, it’s spatial overlap with a mask at the edge of the brain, a mask of cerebrospinal fluid, and finally the proportion of high-frequency signal content. Noise components were detected and removed prior to any temporal filtering so the resulting cleaned data were temporally high-pass filtered using a 0.01Hz filter. Supplemental analyses and figures examining the remaining associations between participant age and common motion metrics (e.g. mean frame-to-frame motion) relevant to functional connectivity measurement are included in Supplemental Material.

The resulting image was resampled into a 2mm isotropic standard space and smoothed to a 5mm isotropic full-width half maximum smoothness using 3dBlurToFWHM in Analysis of Functional NeuroImages (AFNI) software [Cox, 1996]. This was to ensure that all subjects’ data had similar effective spatial smoothness [Scheinost et al., 2014].

#### Adult Data

A gradient echo sequence was collected with a TR/TE of 2500 ms/30 ms, flip angle of 80°, 38 3mm slices (with an additional slice gap of 0.33mm) with a field of view of 216 mm × 216 mm, leading to an in-plane resolution of 3mm. A total of 260 volumes were initially collected but the first 4 were discarded to allow magnetization to reach equilibrium. As with the paediatric fMRI data, each resulting time-series was corrected for inter-scan motion using MCFLIRT and had the middle functional scan rigidly registered to its T1-weighted image. The resulting motion characteristics were calculated as in O’Muircheartaigh et al (2015) and datasets were excluded if there were more than 25 motion spikes of scan-to-scan motion surpassing 0.2mm (as in the paediatric data, this is 10% of the available data). As in the paediatric data, the AROMA pipeline was again employed to remove structured noise components. The resulting denoised timeseries was then resampled into a 2mm isotropic standard space (using the non-linear transformation calculated earlier) and smoothed with a 5mm isotropic Gaussian kernel.

### Amygdala Subregions of Interest

Amygdala subregions were derived from a population atlas of architectonic profiles transformed to a standard population MRI atlas [Amunts et al., 2005] provided with the FSL package. This atlas was thresholded and binarized at 50% probability for the adult analysis. For the paediatric functional analysis, this atlas was first transformed to this study’s paediatric template space and thenthresholded at 50% probability. This threshold of 50% has has been previously used for studies in both adults and young children [e.g. Alarcón et al., 2015; Ball et al., 2007; Gabard-Durnam et al., 2014; Qin et al., 2012; Roy et al., 2009; Roy et al., 2013]. No subregions overlapped in space (see **Figure 1** for an image of the masks rendered on an average T_1_ weighted image and functional MRI image). All subregions of interest were bilateral, given both the high degree of similarity between left and right amygdala functional connectivity maps previously reported in adults (Roy et al. 2009) as well as to facilitate comparison with prior studies in older developmental samples that used bilateral subregions of interest [Gabard-Durnam et al., 2014; Qin et al., 2012]. Supplemental analyses examining differences between the right and left amygdala seeded connectivity did not return any significantly different connections (not reported).

**Figure 1:**
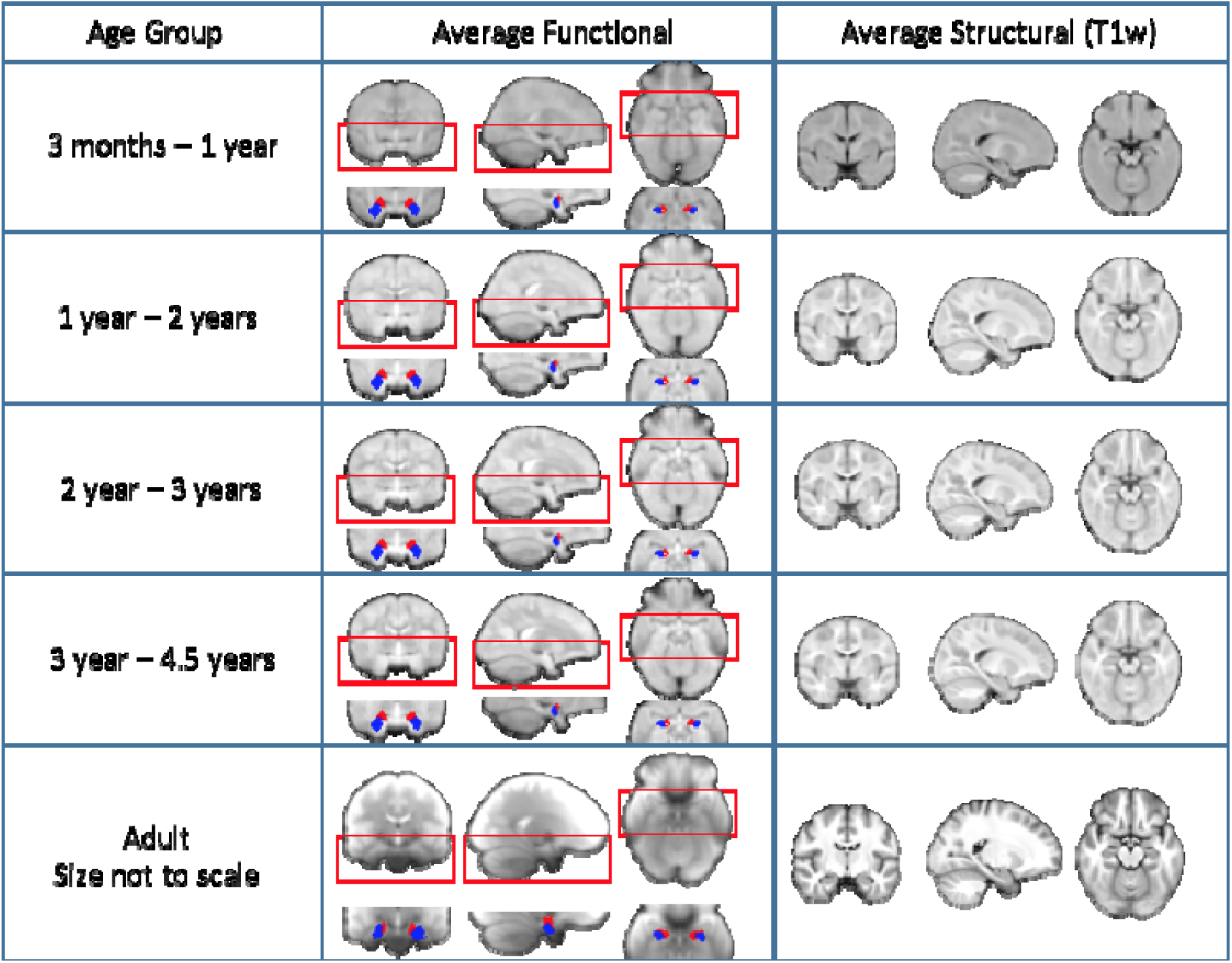
Average functional and structural (T1-weighted) images after spatial registration to a paediatric standard space. Images are shown at 4 different timepoints in the childhood dataset to illustrate registration performance and the placing of the amygdala seeds. Also shown is the average functional and structural image for the adult dataset in MNI space (note that the adult dataset was collected at a different site with different sequences). The bilateral superficial nuclei seeds are shown in red and the bilateral laterobasal seeds are shown in blue.

### Functional Connectivity

For the whole amygdala seed and for each amygdala subregion individually, we calculated the mean time-series for every subject. Due to the proximity of the amygdala to areas of high vascular density, we also created a mask of regions associated with vascularization using a freely downloadable population atlas [Viviani et al., 2009] available here: http://www.uniklinik-ulm.de/struktur/kliniken/psychiatrie-und-psychotherapie/klinik-fuer-psychiatrie-und-psychotherapie-iii-ulm/home/forschung/clinical-neuroimaging/digital-atlas.html. We tested whole-brain functional connectivity with the whole amygdale in a single-subject general linear model (GLM). We tested the two amygdala subregions together in a separate single-subject GLM. These models also included the timeseries of the vascular region as a confound regressor. For each subject, the GLMs were tested using FSL’s Feat.

A cross-sectional GLM was designed to test the average connectivity per contrast, including age, sex, average absolute scan-to-scan motion, average relative scan-to-scan motion as further covariates. Our contrasts of interest were average connectivity and relationships with age and sex. The resulting parameter estimates for the whole amygdala and the amygdala subregion contrasts were then input to a higher-level group analysis using this GLM and tested by permutation using FSL’s Randomise. Multiple comparison correction for each contrast was performed using the cluster extent option in Randomise (thresholding based on the null distribution of cluster size), with an initial cluster forming threshold of t>3.1.

In addition, to explore non-linear age-related changes during this developmental period, two additional sets of GLMs were run as secondary exploratory analyses, where the first set included a quadratic age term, and the second set included a natural log of age term. Both covariates were orthogonalised with respect to the linear age term. Results were corrected for multiple comparisons as before, using the cluster extent option in Randomise, with an initial cluster forming threshold of t>3.1. There were no statistically significant functional connectivity patterns associated with the age-squared non-linear term that survived correction, so the following results are from set of analyses examining only the linear age-related and non-linear natural log(age) effects.

Finally, for the whole amygdala and each amygdala subregion separately, the thresholded average connectivity patterns from the developmental and adult samples were overlaid to compare the spatial overlap of connectivity between samples. Spatially overlapping connectivity with the same valence (e.g. positive connectivity) in the developmental and adult samples (concordant connectivity) was distinguished from overlapping connectivity with differing valence (e.g. one sample has positive connectivity, the other has negative connectivity) between samples (discordant connectivity).

## Results

### Average Functional Connectivity with the Amygdala

In the paediatric sample, the whole amygdala had average positive connectivity with ventral cortical and subcortical structures. Significant positive connectivity with cortical structures was limited to the temporal cortex (middle and inferior gyri), insula, and posterior orbitofrontal cortex (OFC), while connectivity with subcortical structures included the basal ganglia, striatum (ventral and dorsal), and thalamus (Figure 2a). The whole amygdala had concordant overlap with adult positive functional connectivity across all of these regions (Figure 2b and c). In the paediatric sample, the whole amygdala had average negative connectivity with dorsal cortical regions. Significant negative connectivity was observed with visual cortex, precuneus, motor and somatosensory cortex, posterior, middle, and dorsal anterior cingulate cortex (Figure 2a). The whole amygdala had concordant overlap with adult negative functional connectivity across the dorsal and middle cingulate, and the visual cortex (Figure 2b and c). The whole amygdala had discordant overlap across posterior cortical regions including the motor and somatosensory cortex, precuneus, posterior cingulate, and superior temporal gyrus (Figure 2d).

**Figure 2:**
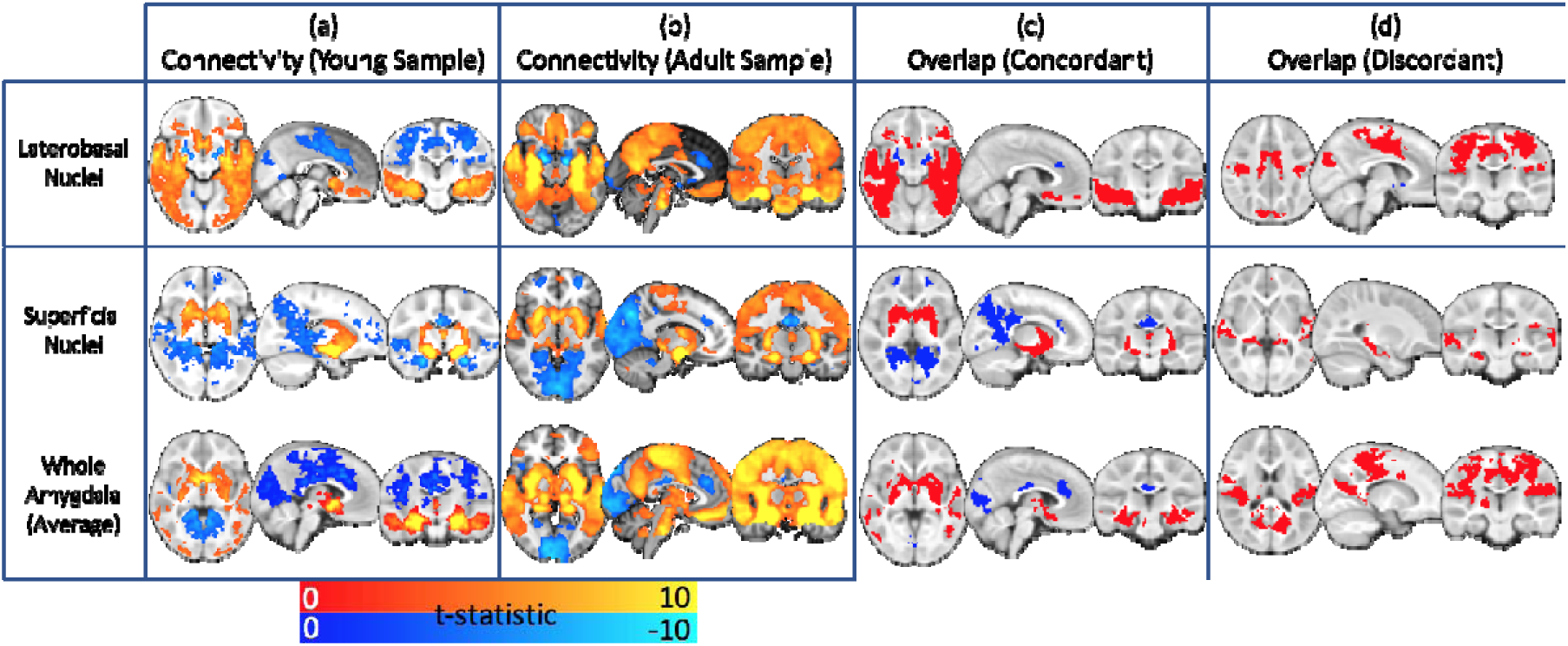
Average functional connectivity with the amygdala. Whole brain functional connectivity at rest between the subdivisions of the amygdala and the rest of the brain (top two rows) and the entire amygdala (bottom row) in the early childhood sample collected as part of this study (column a) and in an adult dataset analysed in the same way (column b). Images are shown after multiple comparison correction (p<0.05, corrected for cluster extent) Column c shows the spatial overlap between them (red is positive connectivity in both samples, blue is negative) and, finally, column d shows regions where connectivity is discordant between age groups, specifically where connectivity is negative in the younger age group and positive in the older age group. Columns c and d are illustrative and are not formal conjunction tests.

The LB subregion had average positive connectivity with frontal and ventral cortical and subcortical regions (Figure 2a). The cortical regions included bilateral OFC (medial and lateral), ventral medial PFC (vmPFC), superior frontal gyrus, motor and somatosensory cortices, temporal cortex (superior, middle, and inferior gyri), precuneus, and visual cortex (cuneus). Subcortical regions included extensive areas of the striatum (ventral striatum and putamen), the hippocampus and parahippocampal gyrus, the thalamus, and the cerebellum. The LB had average negative connectivity with distributed regions including bilateral dmPFC (supplemental motor area), dorsal ACC, pre- and post-central gyri, visual cortex (cuneus and lingual gyri), fusiform gyrus, and dorsal striatum (caudate).

The SF subregion had predominantly average negative connectivity. However, positive connectivity was observed with subcortical structures including bilateral amygdala, thalamus, and striatum (ventral striatum, caudate, putamen) (Figure 2a). The SF had average negative connectivity with bilateral rostral vmPFC (Brodmann Area 11), mPFC (Brodmann Areas 9,10), dorsal cingulate, supplemental motor area, middle frontal gyrus, precentral gyrus, superior and middle temporal gyri, supramarginal gyri, right-hemisphere angular gyrus, precuneus, cuneus, extensive posterior cingulate, fusiform gyrus, hippocampus and parahippocampal gyri, and thalamus.

### Developing Functional Connectivity with the Amygdala

For the whole amygdala and amygdala subregions, associations between connectivity and increasing age were localized to sensory and motor-related regions (Figure 3). The whole amygdala showed decreasing connectivity with age for a region of bilateral thalamus, shifting from positive connectivity at age 3 months to negative connectivity by age 5 years (Figure 3b and c). When broken down to LB and SF connectivity, patterns of age-related change differed between structures.

**Figure 3:**
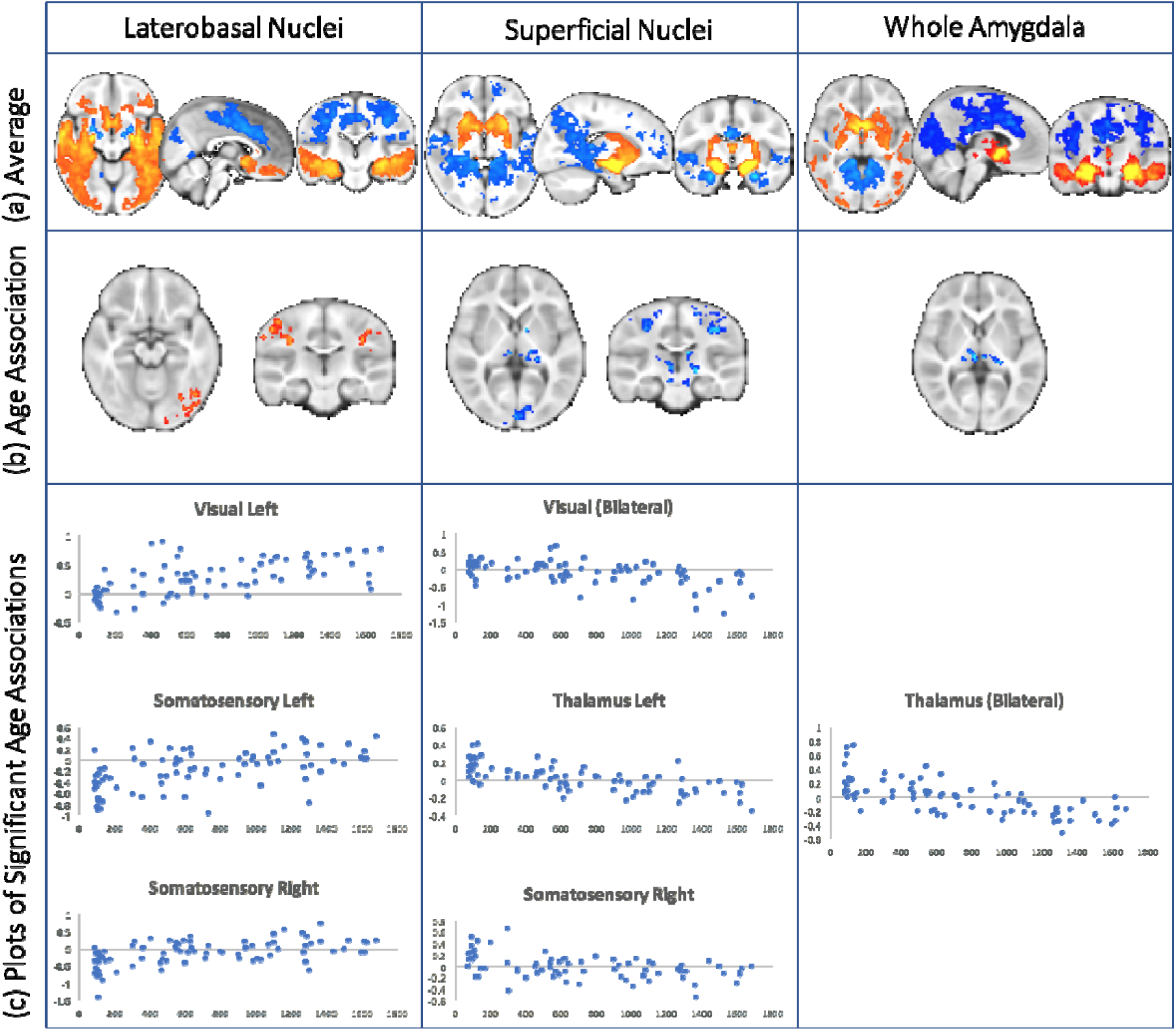
Age-related changes in amygdala functional connectivity. Whole brain functional connectivity at rest between the subdivisions of the amygdala / whole amygdala and the rest of the brain (a) and their association with age middle row (b, hot colours denote positive association and cold negative). The plots (c) illustrate individual differences of connectivity strength (beta, y-axis) with age (in days, x-axis).

The LB subregion showed positive age-related changes in connectivity across sensory and motor cortical regions, reflecting both reductions in negative connectivity and increases in positive connectivity. The LB subregion showed a reduction in negative connectivity with the bilateral postcentral gyrus (somatosensory cortex), shifting from negative connectivity at age 3 months to no significant connectivity by age 5 years (Figure 3b and c). The LB subregion also showed increasing positive connectivity with age for a bilateral occipital cortex region, such that no statistically significant connectivity was observed at age 3 months, and positive connectivity emerged by age 5 years (Figure 3b and c).

In contrast, the SF subregion showed negative age-related changes in connectivity across subcortical and cortical sensory and motor regions. The SF subregion connectivity with the thalamus shifted from positive connectivity at age 3 months to negative connectivity by age 5 years (Figure 3b and c). Similarly, the SF subregion connectivity with the somatosensory cortex shifted from positive connectivity at age 3 months to negative connectivity by age 5 years (Figure 3 b and c). An overlapping SF-somatosensory cortical connection showed nonlinear (natural log) age-related decrease in connectivity during this period as well. The SF subregion also showed increasingly negative connectivity with occipital cortex, such that no statistically significant connectivity was observed at 3 months of age, and negative connectivity emerged by 5 years of age (Figure 3).

## Discussion

Despite the amygdala’s critical and basic role in shaping emotional and social behavior early in life, we currently have little knowledge regarding the human amygdala’s functional development during this period. Here, we characterized the functional (resting-state) networks of the whole amygdala and the amygdala’s LB and SF subregions, and their age-related changes in a large cohort of typically-developing children from 3-months to 5-years of age. We demonstrated that the whole amygdala functional connectivity pattern is already largely present in early childhood, recapitulating the patterns observed in middle childhood and adulthood, with the interesting exception of medial prefrontal cortex connectivity. Similarly, we have shown spatially distinct functional connectivity patterns for each amygdala subregion, suggesting network distinctions between subregions are evident and observable during infancy. We have also shown age-related maturational changes in the whole amygdala and amygdala subregions’ resting-state connectivity limited to sensory and motor-related regions, a pattern unique to this early post-natal developmental period.

Distinct patterns of average connectivity and age-related changes in connectivity separated each of the amygdala subregions in infancy and early childhood, and were consistent with connectivity observed later in development. Overall, the observed pattern of both positive LB connectivity with temporal and ventral cortical regions and negative LB connectivity with dorsal and posterior cortical regions is highly consistent with stable patterns observed later in development and in maturity [Alarcón et al., 2015; Gabard-Durnam et al., 2014; Qin et al., 2012; Roy et al., 2013]. A notable exception to this consistency was the observed LB-PFC connectivity. The LB has been shown to have particularly robust anatomical connections with the vmPFC across species (Aggleton et al., 1980; Ghashghaei and Barbas, 2002; McDonald, 1998), but prior developmental studies have not consistently identified LB-vmpfc or LB-OFC functional connectivity (age-related increases across middle childhood and adolescence in Gabard-Durnam et al., 2014, but present in Qin et al., 2012 in middle childhood, and whole-amygdala OFC/vmPFC connectivity at preschool age reported by Shen et al., 2016). Here we show that LB-OFC and LB-vmPFC connectivity was already present in this sample of infants and young children, suggesting early, coordinated communication does occur between these regions. Additionally, functional connectivity between the LB and limbic regions, such as the vmPFC, striatum, and medial temporal lobe (e.g. hippocampus), has been implicated as a core network for stimulus valuation [Ghashghaei and Barbas, 2002; Pessoa and Adolphs, 2010], fear/threat memory, and conditioned response retrieval [LaBar and Cabeza, 2006; Maren and Quirk, 2004; Sierra-Mercado et al., 2011]. Robust positive connectivity between the LB and all of these regions was already seen in this early developmental sample. A recent report in 1-month olds has shown that amygdala-striatal and amygdala-vmPFC connectivity already correlate with later fear and cognitive development profiles [Graham et al., 2015a]. The pattern of connectivity from our findings would suggest that connectivity with the LB subregion (the largest cluster of nuclei) in particular may be driving these amygdala results [Graham et al., 2015a]. LB-hippocampal-frontal functional connectivity also increases through adolescence, suggesting continued refinement of this circuitry occurs after this period of stable connectivity in infancy and early childhood [Gabard-Durnam et al., 2014; Qin et al., 2012; Silvers et al., 2016].

Age-related changes in LB connectivity were also observed during infancy and early childhood with sensori-motor and occipito-temporal regions. LB-occipitalconnectivity increased with age during infancy and early childhood in parallel with reports of extensive changes in the face-processing capacities subserved in part by amygdala-occipito-temporal connections [Anderson et al., 2016; Cassia et al., 2009; Picozzi et al., 2009; Simion and Giorgio, 2015]. Future work focusing on the emergence of this LB-occipito-temporal functional connectivity and face-processing directly can elucidate whether this early connectivity drives these face-processing developments. The LB also showed refinement of connectivity with the sensorimotor cortex during infancy and early childhood. Specifically, negative connectivity between the LB and several regions within bilateral sensorimotor cortex in infancy was reduced to no significant connectivity between these regions. This selective reduction of LB connectivity with parts of the sensorimotor cortex may reflect spatial refinement of this functional connection during early development. The LB maintained significantly negative connectivity with other regions of sensorimotor cortex throughout this period, though. This finding is consistent with the negative connectivity maintained between the LB and regions of the sensorimotor cortex across childhood and adolescence into adulthood [Gabard-Durnam et al., 2014; Roy et al., 2009]. Moreover, the LB-sensorimotor negative connectivity remaining after infancy has been observed to undergo strengthening (i.e. increasingly negative valence) during the childhood and adolescence periods (Gabard-Durnam et al., 2014). Together, the present and prior results suggest that LB-sensorimotor functional connectivity undergoes a series of refinements during a protracted developmental trajectory. Future studies including behavioral measures of motor development or tasks including motor responses during affective contexts may reveal the behavioral consequences of these different connectivity refinements across development.

SF patterns of functional connectivity and age-related changes in connectivity were also highly consistent with those identified in later development. In particular, the positive functional connections with ventral caudate and anterior putamen regions and negative connectivity with broad posterior-cingulate and superior-frontal regions have all been observed as stable patterns in older populations. The SF’s positive subcortical connectivity is also in line with preliminary studies in adult humans implicating the SF in olfactory and social incentive processing [Bzdok et al., 2013; Goossens et al., 2009; Heimer and Van Hoesen, 2006; Koelsch et al., 2013]. Although positive SF connectivity extending to ventral cortical regions has been observed in older populations, this was not observed in the present sample, and it may be that this connectivity emerges in early childhood (at the junction of sample age-ranges between this study and prior studies) [Bzdok et al., 2013; Gabard-Durnam et al., 2014]. It should be noted that the SF’s spatial proximity to ventricles tempers the extent of interpretation for this region’s connectivity at typical fMRI resolution.

The age-related changes in SF connectivity were also consistent with changes that continue through adolescence in older samples, suggesting contiguous, gradual refinement occurs in the SF-network. Specifically, while there was robust positive SF connectivity with much of the thalamus, consistent with both mature resting-state and stimulus-elicited SF-networks [Bzdok et al., 2013; Koelsch et al., 2013; Roy et al., 2009], there were also regions of the thalamus where connectivity with the SF changed to negative connectivity by early childhood, and thalamic connectivity continues to change (seen as positive connectivity attenuating) through adolescence [Gabard-Durnam et al., 2014]. SF-occipital connectivity was also observed to change such that negative connectivity emerged by early childhood, mirroring the same shift from no SF connectivity to negative connectivity with other regions of the occipital lobe across childhood and adolescence [Gabard-Durnam et al., 2014; Roy et al., 2009]. Lastly, SF-motor cortex connectivity in the left hemisphere changed from positive to negative connectivity by early childhood, foreshadowing a similar shift between SF and the right-hemisphere motor cortex that begins in middle childhood and persists into adulthood, although the mechanisms supporting this asymmetrical developmental timing remain to be explored. Although the interpretation of negative connectivity is unresolved, prior developmental studies of other functional networks have observed developmental shifts from positive to negative connectivity (e.g. Chai et al., 2014). Moreover, preliminary evidence from both empirical studies and simulations of functional network properties suggests that the emergence of negative connectivity reflects complex regulatory interactions between regions and networks, including reciprocal modulation between regions, suppression of one region by the other, and other inhibitory processes (Cabral et al., 2011; Gopinath et al, 2015; Parente and Colosimo, 2016; Parente et al., 2017). Future studies targeting SF function in specific stimuli contexts across infancy and early childhood can probe the nature of this potentially regulatory relationship with motor cortex.

Although networks continue segregating through adolescence, these spatial patterns of connectivity suggest core network components differentiating the subregions are in place early (Qin et al., 2012; Roy et al., 2013; Gabard-Durnam et al., 2014). This set of findings is consistent with evidence that amygdala subregions segregate structurally early in development [Humphrey, 1968; Payne et al., 2010; Ulfig et al., 2003]. Moreover, for areas where both amygdala subregions showed significant connectivity, frequently the two subregions demonstrated differently-valenced coupling (e.g. one subregion showing positive connectivity, the other showing negative connectivity), suggesting the subregions show functional differentiation from each other for spatially-overlapping connections. For example, the putamen had robust connectivity with the LB and SF subregions, but this connectivity was negative with the LB and positive for the SF. In addition, subregions demonstrated distinct patterns of change in their network connectivity during this period. The LB showed switches to positive connectivity with sensori-motor regions, while the SF showed switches to negative connectivity by early childhood that continue through adolescence. These results suggest the subregions may have disparate trajectories of connectivity development in terms of both timing and network composition beginning in infancy.

Several patterns emerged across the subregions’ connectivity in infancy and early childhood distinguishing this period from amygdala networks in later developmental periods. First, the period of infancy is unique in that amygdala reactivity is absent to stimuli during this time that elicit responsiveness in childhood, but we and others have observed largely intact functional connectivity during infancy [Blasi et al., 2011; Goksan et al., 2015; Graham et al., 2013; Graham et al., 2015a; Qiu et al., 2015]. Thus, future work may examine whether this early established connectivity facilitates the emergence of later amygdala reactivity by childhood.

Secondly, no age-related changes were observed with any subregion and association cortex during this early post-natal period, contrary to later childhood and adolescence (Tottenham and Gabard-Durnam, 2017). Together with previous studies, these results suggest that amygdala-subcortical and sensory-cortex connectivity begins refinement prior to childhood and continues through adolescence, while connectivity changes with associative and frontal cortical areas may begin after early childhood [Gabard-Durnam et al., 2014; Qin et al., 2012; Roy et al., 2013].

In particular, positive connectivity of the whole amygdala or either subregion was not observed during infancy and early childhood with the perigenual medial anterior cingulate. This finding stands in contrast to a recent report of amygdala resting-state connectivity in early childhood (Shen et al., 2016), although differences in how the amygdala seeds were defined between the samples may in part account for these differences. Moreover, the present finding is consistent with another well-powered report in middle childhood (Thijssen et al., 2017) and with prior cross-sectional and longitudinal reports that positive connectivity between the amygdala and this perigenual mPFC region underlying emotion regulation emerges in late childhood and adolescence (Qin et al., 2012; Gabard-Durnam et al., 2014; Alarcón et al., 2015; Gabard-Durnam, Gee, et al., 2016; Tottenham and Gabard-Durnam, 2017). Aside from differences in data processing approaches across studies, given Thijssen and colleagues’’s finding that amygdala-perigenual mPFC connectivity differs as a function of parent-child dynamics, it is also possible that youth- and family-level differences across these developmental samples (e.g. youth internalizing levels, attachment styles, parental stress levels) have contributed to the disparate findings across study sites. Future studies may further examine how such factors contribute to amygdala-perigenual mPFC connectivity during development to clarify the prior body of work.

Several caveats should be noted in considering these results. First, dramatic changes in brain size occur during this early developmental period [Huttenlocher and Dabholkar, 1997; Knickmeyer et al., 2008]. To address the possible variability in tissue partial volume in different regions, we matched spatial smoothness across subjects [Scheinost et al., 2014]. Nonetheless, addressing differential head size in developmental studies remains challenging as a simple covariate for head size (for example) is highly collinear with age. In fMRI generally, and in developmental fMRI in particular, motion has also emerged as a serious potential confound for connectivity studies [e.g. Hallquist et al., 2013; Power et al., 2011; Satterthwaite et al., 2012; Scheinost et al., 2014], so this study employed very strict motion criteria per current recommendations. However, there is no way presently to index with certainty that no remaining motion-related artifact exists [Power et al., 2015]. Our use of the architectonic subregion maps in infants is also novel, but untested. However, the overlap in findings with studies with youths as young as 4-years through adulthood is consistent with these maps delineating the same subregions across development [Alarcón et al., 2015; Gabard-Durnam et al., 2014; Kim et al., 2011; Qin et al., 2012; Roy et al., 2009; Roy et al., 2013]. These consistencies also suggest that the scanning state (asleep in this study, awake in other studies) may not alter amygdala resting-state connectivity significantly, consistent with prior reports of resting-state connectivity patterns persisting in sleep states [Fukunaga et al., 2006; Graham et al., 2015b; Horovitz et al., 2008; Larson-Prior et al., 2009, Shen et al, 2016], although verification with further study is needed. The amygdala is also adjacent to ventricles and dense vasculature. This means that, at typical fMRI resolution and sampling rate, there is a risk that the timeseries is confounded by physiological noise [Boubela et al., 2015; Lund et al., 2006; Murphy et al., 2013]. We used a data-driven technique to identify and remove structured noise from the fMRI data but, given the concerns about venous signal in particular [Boubela et al., 2015], we also included signal from a population atlas of vascular density [Viviani et al., 2009]. It should also be noted that the acquisition time for the resting-state scans in the present study was fairly short (~5 minutes). Although resting-state connectivity has been shown to stabilize as rapidly as within 5 minutes (Van Dijk et al., 2010), it is possible that the functional connectivity patterns observed would undergo further changes or stabilization with longer measurements. Lastly, longitudinal studies, that can map out the developmental trajectories of these subregions within individuals, are needed to confirm the patterns currently observed.

Cumulatively, these findings illustrate that the amygdala’s LB and SF subregions have functional circuits in place by infancy, and progressive shaping of motor and sensory circuit components occurs during this period. Subregions can be distinguished from each other by connectivity patterns as well as their age-related changes in connectivity. Indeed, these results suggest that the subregions may have disparate trajectories of network construction across development requiring future longitudinal study. Moreover, the robust amygdala connectivity observed in infancy seems to temporally precede observations of reactivity. These findings present the first information about typical age-related changes in human amygdala functional network development prior to childhood. Together, these results represent an important initial step in understanding the early development of amygdala networks and their dynamics, central to sculpting emotional and social behavior.

## Acknowledgements

The authors declare no competing financial interests. The authors wish to thank all the families that donated their time to take part in this research.

## Supplemental Material

**Supplemental Figure 1.**
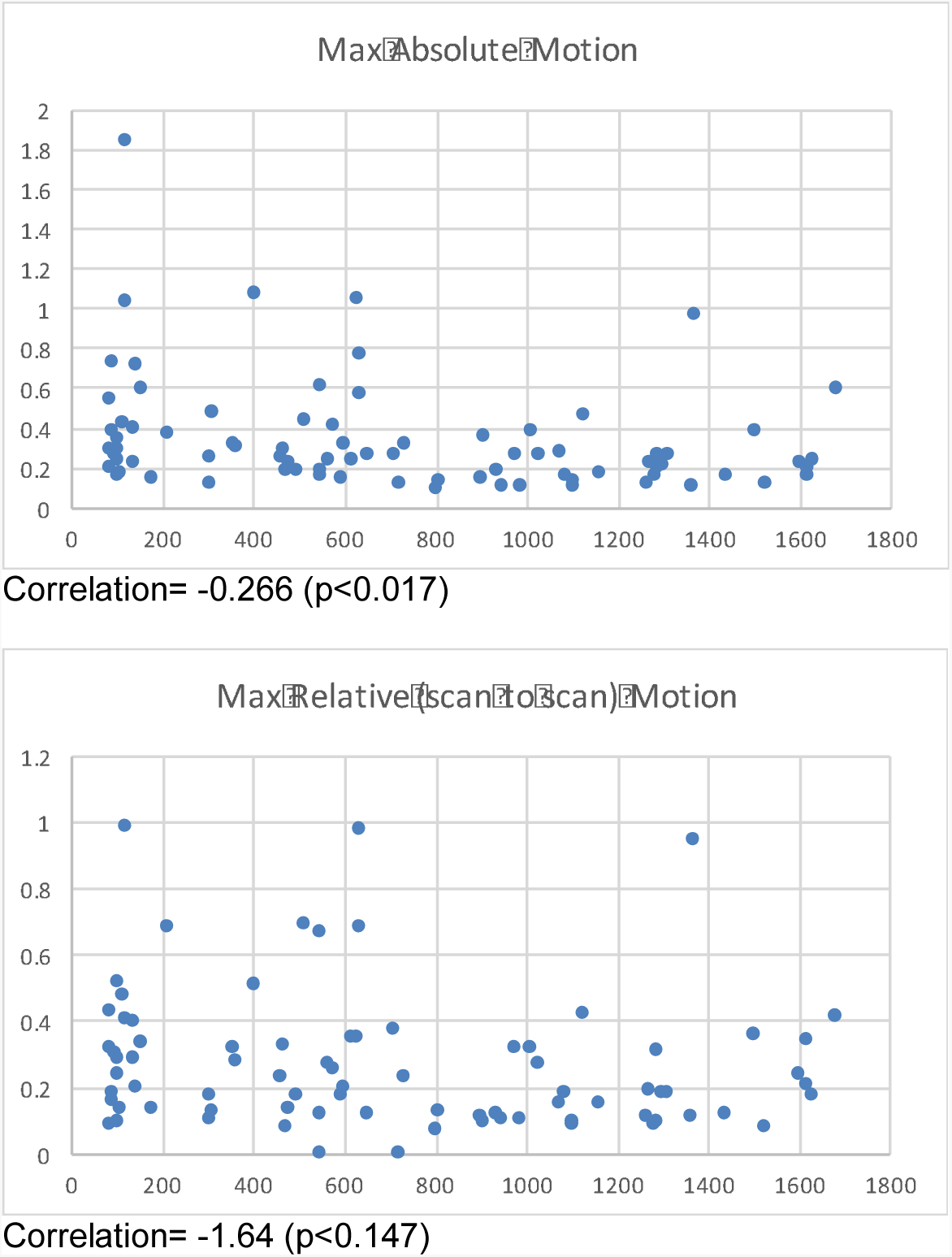
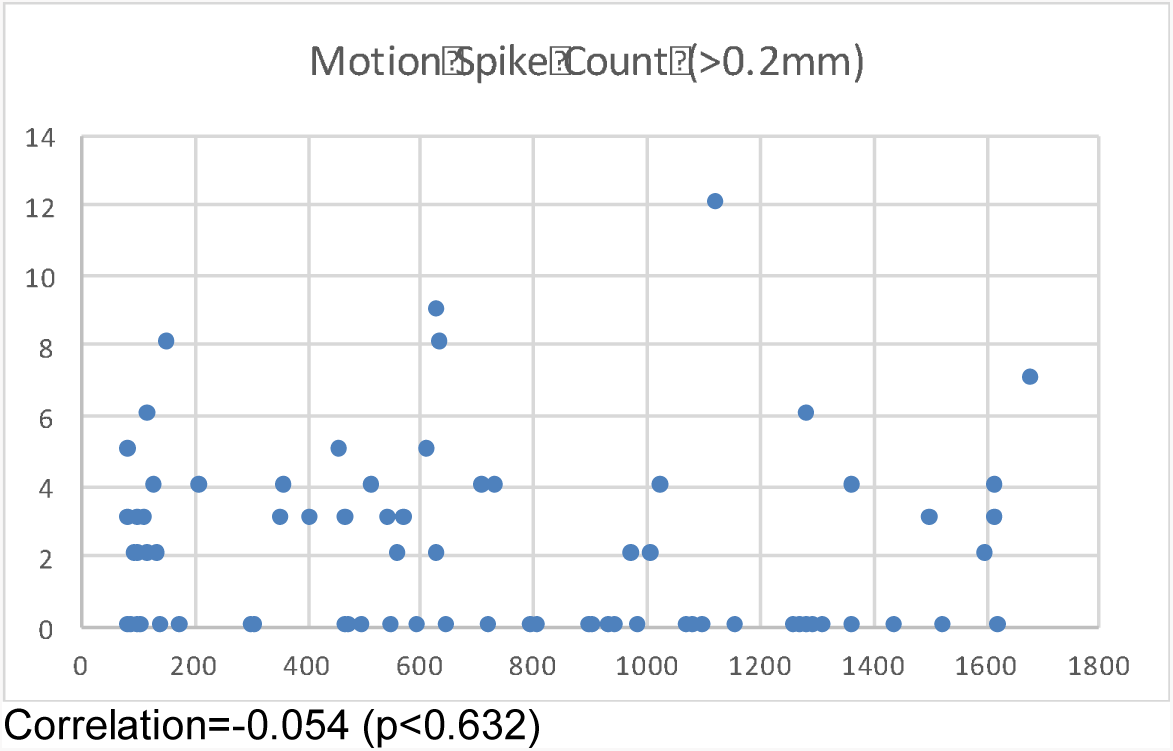
Associations between participant age and common motion metrics calculated from the timeseries used in the main manuscript’s analyses. Pearson correlation tests examining the motion metric’s association with age are reported below the corresponding plot.

